# Top-down beta enhances bottom-up gamma

**DOI:** 10.1101/054288

**Authors:** Craig G. Richter, William H. Thompson, Conrado A. Bosman, Pascal Fries

**Author notes:** Correspondence should be addressed to Dr. Craig Richter or Dr. Pascal Fries, Ernst Strüngmann Institute (ESI) for Neuroscience in Cooperation with Max Planck Society, Deutschordenstraβe 46, 60528 Frankfurt, Germany.,. C.R. and W.T. contributed equally to this study.

## Abstract

**Abstract:** Several recent studies have demonstrated that the bottom-up signaling of a visual stimulus is subserved by interareal gamma-band synchronization, whereas top-down influences are mediated by alpha-beta band synchronization. These processes may implement top-down control of stimulus processing if top-down and bottom-up mediating rhythms are coupled via cross-frequency interaction. To test this possibility, we investigated Granger-causal influences among awake male macaque primary visual area V1, higher visual area V4 and parietal control area 7a during attentional task performance. Top-down 7a-to-V1 beta-band influences enhanced visually driven V1-to-V4 gamma-band influences. This enhancement was spatially specific and largest when beta-band activity preceded gamma-band activity by ∼0.1 s, suggesting a causal effect of top-down processes on bottom-up processes. We propose that this cross-frequency interaction mechanistically subserves the attentional control of stimulus selection.

**Significance Statement:** Contemporary research indicates that the alpha-beta frequency band underlies top-down control, while the gamma-band mediates bottom-up stimulus processing. This arrangement inspires an attractive hypothesis, which posits that top-down beta-band influences directly modulate bottom-up gamma band influences via cross-frequency interaction. We evaluate this hypothesis determining that beta-band top-down influences from parietal area 7a to visual area V1 are correlated with bottom-up gamma frequency oscillations from V1 to area V4, in a spatially specific manner, and that this correlation is maximal when top-down activity precedes bottom-up activity. These results show that for top-down processes such as spatial attention, elevated top-down beta-band influences directly enhance feedforward stimulus induced gamma-band processing, leading to enhancement of the selected stimulus.

## Introduction

Many cognitive effects in vision can only be explained by invoking the concept of top-down influences (Gilbert and Sigman, 2007). For example, when top-down influences pre-allocate attention to specific spatial locations, stimulus processing is more accurate and/or faster. Correspondingly, neurons in higher visual areas show enhanced firing when processing attended stimuli. These neurophysiological consequences of top-down control must be mediated by corresponding anatomical projections. Indeed, anatomical studies have documented projections in the top-down direction that are as numerous as bottom-up projections. Bottom-up and top-down projections show different characteristic laminar patterns of origin and termination, and the pattern of interareal pairwise projections abides by a global hierarchy (Felleman and Van Essen, 1991; Hilgetag et al., 1996; Markov et al., 2014). Recently, it has been shown in both macaque and human visual cortex, that the pattern of anatomical projections is closely correlated to a pattern of frequency-specific directed interareal influences. Influences mediated by bottom-up projections are primarily carried by gamma-band synchronization; influences mediated by top-down projections are primarily carried by alpha-beta-band synchronization (Bastos et al., 2012; Bosman et al., 2012; Grothe et al., 2012; Jia et al., 2013; van Kerkoerle et al., 2014; Bastos et al., 2015b; 2015c; Michalareas et al., 2016). A similar association of higher frequency bands with bottom-up and lower frequency bands with top-down signaling has also been found in other systems (Colgin et al., 2009; Bieri et al., 2014; Fontolan et al., 2014) and related to respective task demands (von Stein and Sarnthein, 2000; Buschman and Miller, 2007).

Here, we investigate directly whether top-down influences actually modulate bottom-up influences. We simultaneously assess influences in both directions through multi-area electrophysiology and frequency-resolved Granger causality analysis. We find that on an epoch-by-epoch basis, top-down beta-band influences enhance bottom-up gamma-band influences. This effect is spatially specific, i.e. bottom-up gamma-band influences depend most strongly on the top-down beta-band influences that are directed to the origin of the bottom-up influence. Spontaneous enhancements in top-down beta-band influences are followed ∼0.1 s later by enhancements in bottom-up gamma-band influences, suggestive of a causal relation.

We suggest that this is a mechanism for the implementation of spatially selective attention. Attentional control areas in posterior parietal cortex contain maps of visual space, with neuronal ensemble activity describing the spatial location of a subject’s attention (Bisley and Goldberg, 2003). These neurons likely exert top-down beta-band influences on retinotopically corresponding parts of early visual cortex, which in turn enhance the bottom-up forwarding of visual stimuli.

## Materials and Methods

### Visual stimulation and behavioral task

The experiment was approved by the ethics committee of the Radboud University Nijmegen (Nijmegen, The Netherlands). Two adult male macaque monkeys (monkey K and monkey P, both macaca mulatta) were used in this study. During experiments, monkeys were placed in a dimly lit booth facing a CRT monitor (120 Hz non-interlaced). When they touched a bar, a fixation point was presented, and gaze had to remain within the fixation window throughout the trial (monkey K: 0.85 deg radius, monkey P: 1 deg radius), otherwise the trial would be terminated and a new trial would commence. Once central fixation had been achieved and a subsequent 0.8 s pre-stimulus interval had elapsed, two isoluminant and isoeccentric drifting sinusoidal gratings were presented, one in each visual hemifield (diameter: 3 deg, spatial frequency: ≈ 1 cycle/deg, drift velocity: ≈1 deg/s, resulting temporal frequency: ≈1 cycle/s, contrast: 100‥). Blue and yellow tints were randomly assigned to each of the gratings on each trial (Fig. 1A). Following a random delay interval (monkey K : 1 - 1.5 s; monkey P : 0.8 - 1.3 s), the central fixation point changed color to match one of the drifting gratings, indicating that this grating was the target stimulus, i.e. the fixation point color was the attentional cue. When the target stimulus was positioned in the visual hemifield contralateral to the recorded hemisphere, we refer to this condition as attend-contra, whereas when the target was in the ipsilateral hemifield with respect to the ECoG grid, this condition is labeled attend-ipsi. Either the target or distracter stimulus could undergo a subtle change in shape consisting of a transient bending of the bars of the grating (0.15 s duration of the full bending cycle). This change could occur at any monitor refresh from 0.75 s to 5 s (monkey K), and 4 s (monkey P) after stimulus onset. Bar releases within 0.15 - 0.5 s after target changes were rewarded. If stimulus changes occurred before the cue indicated which stimulus was the target, reports were rewarded in a random half of trials. Bar releases after distracter changes terminated the trial without reward. Trials were pooled from both contra and ipsi conditions, except where explicit comparisons of these conditions were made.

**Figure 1.**
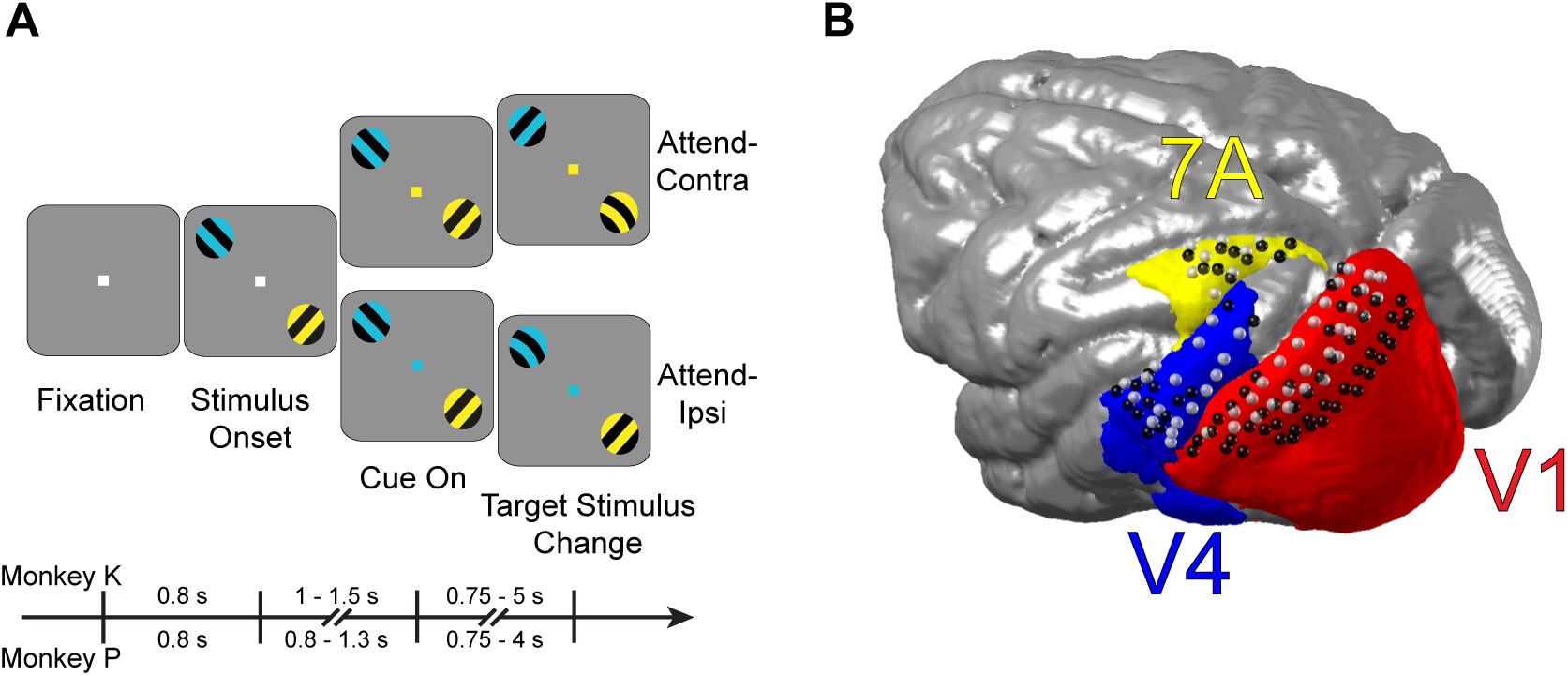
Behavioral task and recording locations. ***A***, The task commenced with a fixation period followed by presentation of two differently colored stimuli. The fixation point color then indicated the visual stimulus to covertly attend in either the visual hemifield ipsilateral (attend-ipsi) or contralateral (attend-contra) to the recording grid. The presentation timings for each monkey are shown as a timeline. ***B***, Recording sites for areas V1 and V2 (red), V4 (blue), and 7A (yellow) from monkey K (light gray spheres) and monkey P (black spheres), co-registered to a common macaque template.

### Neurophysiological recordings and signal preprocessing

LFP recordings were made via a 252 channel electrocorticographic grid (ECoG) implanted subdurally over the left hemisphere (Rubehn et al., 2009). Data from the same animals, partly overlapping with the data used here, have been used in several previous studies (Bosman et al., 2012; Pinotsis et al., 2014; Brunet et al., 2014a; 2014b; 2015; Richter et al., 2015; Vinck et al., 2015; Bastos et al., 2015b; 2015c; Lewis et al., 2016). Recordings were sampled at approximately 32 kHz with a passband of 0.159 – 8000 Hz using a Neuralynx Digital Lynx system. The raw recordings were low-pass filtered at 250 Hz, and downsampled to 1 kHz. The electrodes were distributed over eight 32-channel headstages, and referenced against a silver wire implanted onto the dura overlying the opposite hemisphere. The electrodes were re-referenced via a bipolar scheme to achieve 1) greater signal localization 2) cancellation of the common reference, which could corrupt the validity of connectivity metrics, 3) to reject headstage specific noise. The bipolar derivation scheme subtracted the recordings from neighboring electrodes (spaced 2.5 mm) that shared a headstage, resulting in 218 bipolar derivations, henceforth referred to as “sites” (see Bastos et al. (2015c) for a detailed description of the re-referencing procedure). The site locations are shown as spheres in Figure 1b (monkey K: light gray, monkey P: black).

All signal processing was conducted in MATLAB (MathWorks, USA) and using the FieldTrip toolbox (http://www.fieldtriptoolbox.org/) (Oostenveld et al., 2011). Raw data were cleaned of line noise via the subtraction of 50, 100, and 150 Hz components fit to the data using a discrete Fourier transform. Following trial epoching (see below for details), epochs for each site were de-meaned by subtracting the mean over all time points in the epoch. Epochs exceeding 5 standard deviations of all data from the same site in the same session were rejected. In addition, epochs were manually inspected and epochs with artifacts were rejected. The remaining epochs were normalized by the standard deviation across all data in all epochs from the same site in the same recording session. Subsequently, all epochs of a given site were combined across sessions.

### Region of interest definition

Three regions-of-interest (ROIs) were selected for the current study: V1, V4, and area 7A (referred to simply as “7A”). ROIs were defined based on comparison of the electrode locations (co-registered to each monkey’s anatomical MRI and warped to the F99 template brain in CARET (Van Essen, 2012), with multiple cortical atlases of the macaque (see Bastos et al. (2015c) for a detailed description). Recording sites composing each ROI were co-registered to a common template (INIA19, (Rohlfing et al., 2012)), as were the Paxinos ROI definitions (Paxinos et al., 1999). The V1/V2 combined definition of Paxinos et al. (1999), is shown in Figures 1B, 3G (red) for simplicity due to uncertainty across atlases of the V1/V2 border. Recording site selection was based on multiple atlases with no recording sites selected that were believed to belong to area V2. Based on these ROI definitions, 77 recording sites were selected from area V1 (monkey K: 29, monkey P: 48), 31 from area V4 (monkey K: 17, monkey P: 14), and 18 from area 7A (monkey K: 8, monkey P: 10).

### Segmenting data into periods and epochs

Each successfully completed trial contained three periods: The fixation, the stimulation and the attention period. The fixation period was the time between fixation onset and stimulus onset. During the fixation period, monkeys fixated on a fixation point on a gray screen, and there was no stimulus presented and no cue had been nor was presented during that time. The stimulation period was the time between onset of the stimuli and either a change in one of the stimuli or cue onset. During the stimulation period, monkeys kept fixation, the stimuli were continuously present, one tinted yellow the other blue, chosen randomly, and the fixation point had not yet assumed a color, and thereby the attentional cue had not been given. The attention period was the time between cue onset and a target or distracter change, whichever occurred first. During the attention period, monkeys kept fixation, the stimuli were continuously present with their tints, and the fixation point was tinted in one of these colors, thereby providing the attentional cue.

The fixation, stimulation and attention periods all were of variable lengths across trials. The spectral analysis was based on epochs of fixed lengths. Therefore, the task periods were cut into epochs. Longer trials contributed correspondingly more epochs. We aimed at excluding data soon after events, like stimulus onset and cue onset, to minimize effects of post-event transients and non-stationarities on the metrics of rhythmicity and synchronization. Therefore, from the fixation period, we used the last 0.5 s before stimulus onset; from the stimulation period, the first 0.3 s after stimulus onset were discarded; for the attention period, the first 0.3 s after cue onset were discarded.

For the analyses of Figure 2 and Figure 3, the remaining parts of the described periods were segmented into 0.5 s epochs with 60% overlap. This overlap allows for the application of Welch’s method (Welch, 1967) and was selected as an optimal overlap for the multitaper method, while maintaining a reasonable computational load (Thomson, 1977; Percival and Walden, 1993). This resulted in 6822 fixation epochs (Monkey K: 3384, Monkey P: 3438), 13675 stimulation epochs (Monkey K: 8109, Monkey P: 5566), and a total of 16212 attend epochs (Monkey K: 7275, Monkey P: 8937), of which 8313 were attend-contra epochs (Monkey K: 3819, Monkey P: 4494) and 7899 were attend-ipsi epochs (Monkey K: 3456, Monkey P: 4443).

**Figure 2.**
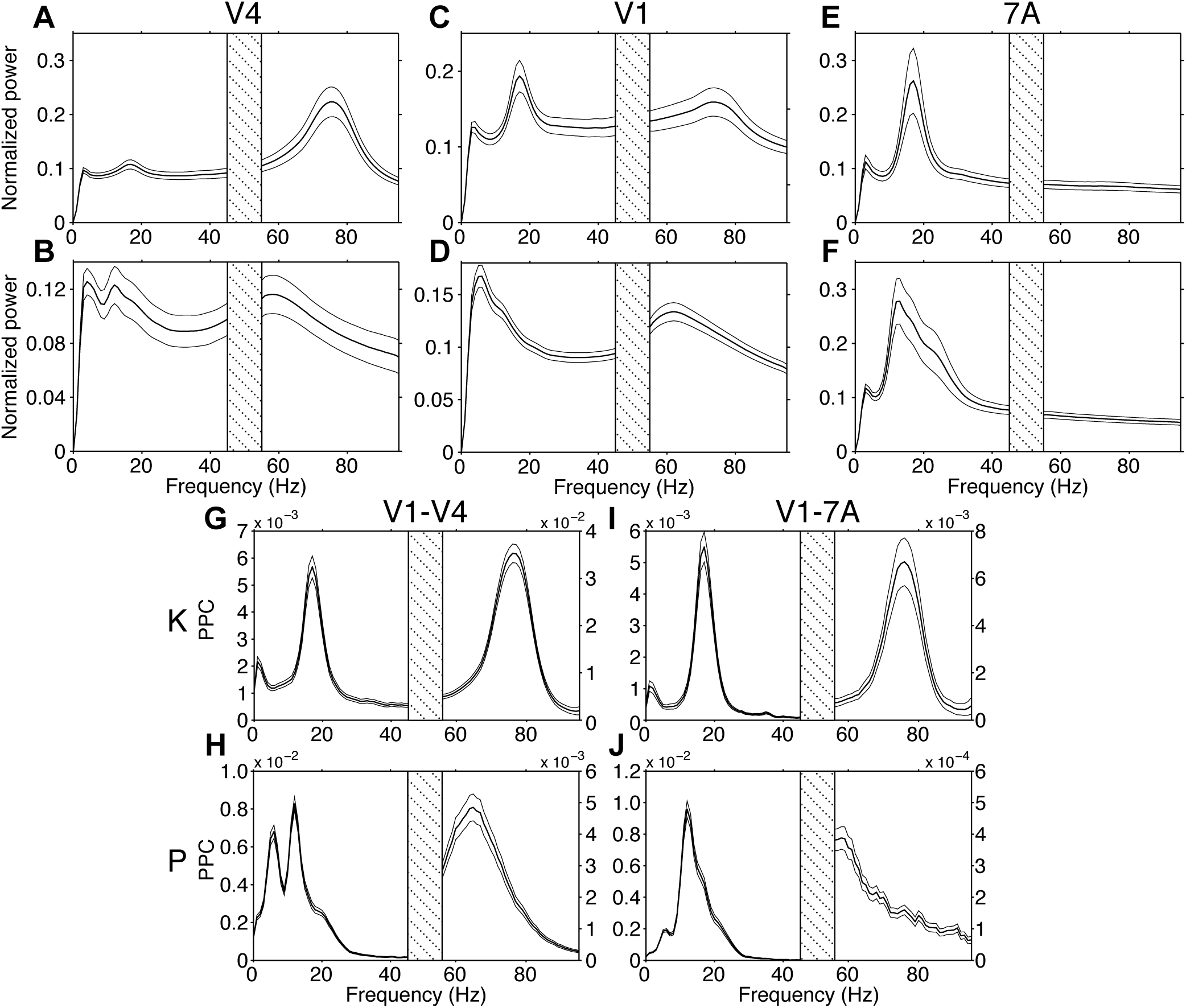
Power spectra and phase locking (PPC) spectra during visual stimulation. V4 (***A***, ***B***), V1 (***C***, ***D***), and 7A (***E***, ***F***) power spectra averaged over all respective site pairs of monkey K (***A***, ***C***, ***E***) and P (***B***, ***D***, ***F***). Power values at each frequency were multiplied by that frequency value to reduce the 1/f component. (***G***-***J***) LFP-LFP PPC for V1-V4 (***G***, ***H***) and V1-7A (***I***, ***J***), for monkey K (***G***, ***I***) and P (***H***, ***J***), respectively. Error regions show ±1 standard error of the mean (SEM) over sites or site pairs. Frequencies from 45-55 Hz were omitted due to line-noise pollution.

For the analyses of Figures 4-7, the same procedure was followed, but using only the attention period and segmenting it into epochs without overlap, because the correlation analyses shown in these figures require independent epochs. This resulted in fewer epochs, totaling 6414 attend epochs (Monkey K: 2655, Monkey P: 3759).

For the analyses of Figure 8, the attention period was segmented into many overlapping 0.5 s epochs, with epochs centers stepped by 0.005 s and epochs fully contained within 0.3-2 s after cue onset. This was done, because the analysis required GC influence time series. It resulted in a total of 878 time series (Monkey K: 398, Monkey P: 480).

### Spectral Analysis of power, phase locking and directed influences

The 0.5 s epochs were tapered and Fourier transformed using the Fast Fourier Transform (FFT). For frequencies from 0-50 Hz, a Hann taper was utilized, whereas for frequencies above 50 Hz, the multitaper method (MTM) was used to improve the spectral concentration of the gamma rhythm (Thomson, 1982; Percival and Walden, 1993). We applied 5 tapers, resulting in a spectral smoothing of +/- 6 Hz. All epochs were zero-padded to 1 s, resulting in a spectral resolution of 1 Hz. The coefficients resulting from the FFT were used to determine power-spectral densities (PSDs) and cross-spectral densities (CSDs), which are the basis for the two employed connectivity metrics: pairwise phase consistency (PPC) (Vinck et al., 2010), and Granger causality (GC) (Granger, 1969; Bressler and Seth, 2011). For the display of power spectra, the power dropoff with frequency, often referred to as 1/f, was partly compensated by multiplying power values at each frequency by the respective frequency values; for example, the power at 40 Hz was multiplied by 40 (Sirota et al., 2008) (Fig. 2A-F). The PPC metric (Fig. 2G-J, Fig. 3A, B), in contrast to the classic coherence metric, does not contain a sample-size bias, that is, it can be directly compared between different numbers of epochs. The GC metric (Fig. 3C-F) was computed directly from the CSDs, using non-parametric spectral matrix factorization, in contrast to the traditional parametric method based on autoregressive modeling (Dhamala et al., 2008). Connectivity metrics were computed for all interareal site pairs of the ROI pairs V1-V4 and V1-7A.

**Figure 3.**
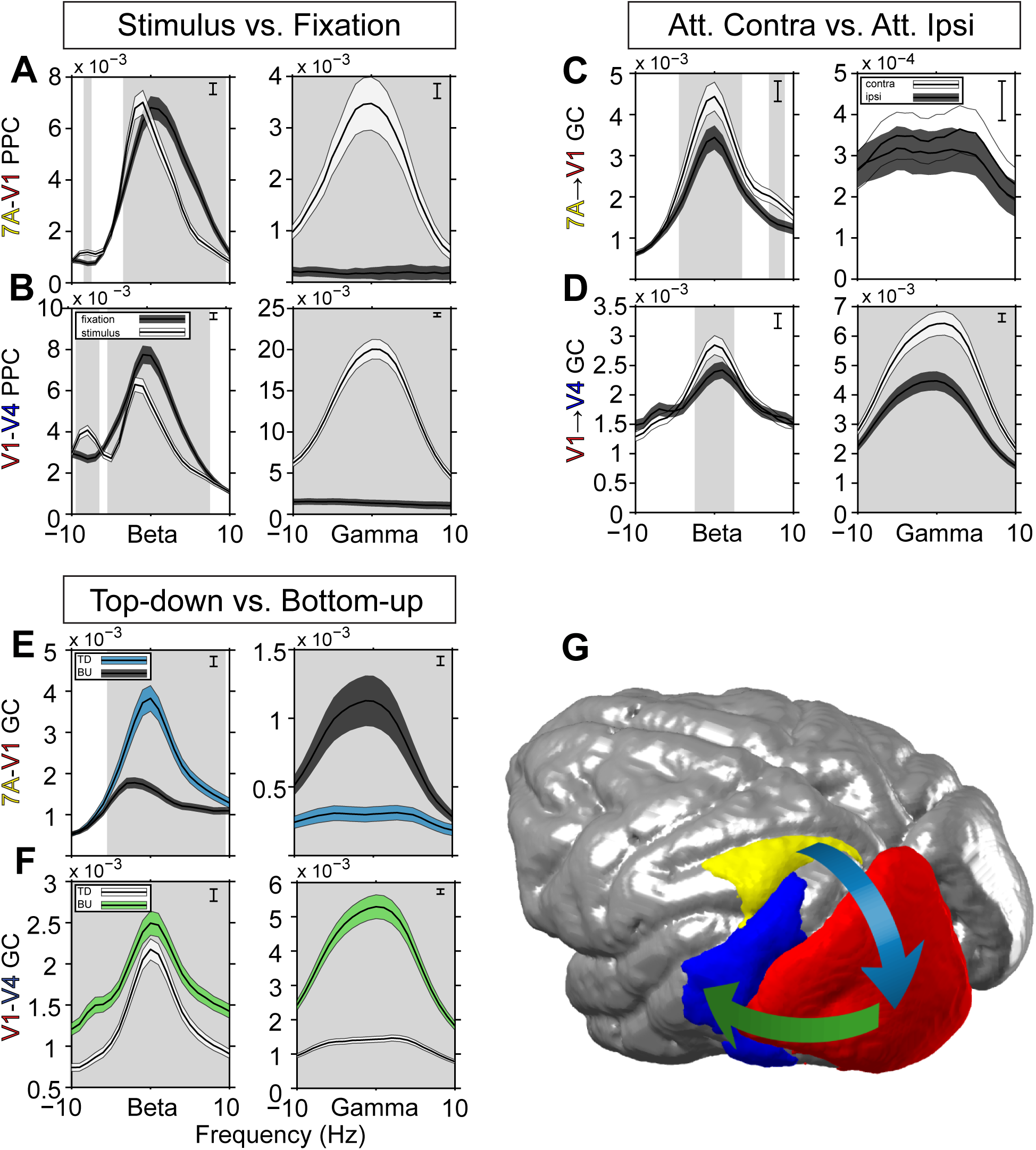
Average phase locking (PPC) and Granger causality (GC) spectra. All spectra were first averaged over site pairs, then over monkeys after alignment to their beta and gamma peak frequencies. ***A***, LFP-LFP PPC for 7A-V1 and ***B***, V1-V4 during fixation (dark shading) and visual stimulation (light shading). Error regions show ±1 standard error of the mean (SEM) over site pairs, merely for illustration. Gray background shading indicates frequency regions with significant condition differences (p<0.05, two-tailed non-parametric randomization test, corrected for comparisons across multiple frequencies). Inset brackets denote the minimum separation required for significance. ***C***, 7A-to-V1 GC influence spectra and ***D***, V1-to-V4 GC influence spectra, for attend-contra (light shading) and attend-ipsi (dark shading). Same statistics conventions as ***A*** and ***B***. ***E***, Bidirectional GC influences for 7A-V1 and ***F***, V1-V4. For 7A-V1, the 7A-to-V1 influences are shown in blue shading, and the V1-to-7A influences in dark shading. For V1-V4, V1-to-V4 influences are shown in green shading, and V4-toV1 influences in light shading. Same statistics conventions as ***A*** and ***B***, but with statistical tests based on a non-parametric bootstrap procedure. ***G***, Top-down modulatory stream (blue arrow) from 7A-to-V1, and bottom-up feedforward path from V1-to-V4 (green arrow).

### Statistical comparison of spectral metrics

Statistical comparison of interaction metrics between conditions used non-parametric randomization, entailing correction for multiple comparisons across frequencies (Maris and Oostenveld, 2007). We illustrate this for PPC. The observed PPC spectra (without randomization) are derived by 1) Calculating PPC spectra across all epochs of a given condition in a given animal, separately for all site pairs of the relevant ROI pair, 2) Averaging PPC spectra across those site pairs, 3) Averaging across the two animals after aligning the peaks of the respective frequency bands. We performed 1000 randomizations, each constituting the null hypothesis. For each randomization, the following steps were performed: 1) Epochs were randomly distributed between conditions. 2) The same steps as for the observed PPC spectra were followed. 3) The maximal absolute difference value across all frequencies was retained and placed into the randomization distribution. The observed differences were compared against the distribution of maximal absolute differences. Observed absolute difference values greater than the 97.5^th^ percentile of the randomization distribution were considered significant at p<0.05. The selection of the maximal absolute difference value per randomization implements the correction for multiple comparisons across frequencies (Nichols and Holmes, 2002). This procedure was used for both PPC and GC influences spectra, separately for the respective frequency bands, either comparing stimulation to fixation or attend-contra to attend-ipsi.

The GC influence metric is known to be biased by sample size (Bastos and Schoffelen, 2015a), thus the number of epochs per attention condition needed to be balanced for each monkey. This was accomplished by finding the condition with the fewest epochs, and randomly selecting this number of epochs from the other condition.

The comparison of top-down versus bottom-up GC was performed on the pooled data from the attend-contra and attend-ipsi conditions. Since this was a within-condition comparison, no balancing of epoch numbers was needed, and all epochs from both attention conditions were used. The statistical analysis of the difference between top-down and bottom-up GC could not be obtained using a non-parametric randomization framework, because top-down and bottom-up GC are not properties of specific sets of epochs, but rather are expressed by all trials simultaneously. Therefore, an alternative statistical approach was used, namely the bootstrap (Efron and Tibshirani, 1994). Like with the randomization approach, the statistic of interest (in this case the top-down/bottom-up GC difference) is recomputed on each bootstrap resample, giving rise to a distribution of surrogate values. Following Efron and Tibshirani (1994), a confidence interval was constructed from the surrogate distribution. To assess the statistical significance at p<0.05 (two-tailed), we find the 2.5^th^ and 97.5^th^ percentile values from the surrogate distribution of differences between top-down and bottom-up GC. This naturally forms the 95% confidence interval such that if zero lies outside of this interval, we may conclude that the result is significant at a level of p<0.05. This method does not control for multiple comparisons, but it can be easily modified to do so using the same logic as employed by Maris and Oostenveld (2007). We performed 1000 bootstrap resamples. For each resample we determined the absolute difference across frequencies between the bootstrap resample spectrum and the average of all bootstrap resamples, and retained the maximum of this value across frequencies. Thus, we are guaranteed to form the largest confidence interval possible across frequencies and in so doing construct an omnibus confidence interval that controls for the multiple comparisons. This confidence interval is applied to each frequency, and where it does not contain zero, the result is significant at p<0.05. To conduct group level statistics, the omnibus statistic is derived from the mean of each bootstrap resample of the difference between top-down and bottom-up spectra across both monkeys (first averaged within-subject across pairs), such that the mean of the empirical difference across the monkeys can be assessed for significance.

### Median split analysis

For the median split analysis, we used a jackknife approach, which leaves out one epoch at a time. The remaining epochs form a jackknife replication (JR). There are as many JRs as there are epochs. 7A-to-V1 beta-band GC (beta peak ±1 Hz) was computed for all JRs. The resulting values were median split, and V1-to-V4 GC computed for the corresponding sets of epochs. The jackknife procedure inverts the distribution of single-epoch GC values (Richter et al., 2015), such that the upper half of JR values corresponds to the lower half of the true single epoch values. The observed differences in V1-to-V4 GC were tested for significance as described for the spectral metrics, using 200 randomizations. The analysis always included a 7A-to-V1 site pair and a V1-to-V4 site pair, sharing the same V1 site. We refer to this configuration as a triplet, or sometimes as 7A-to-V1-to-V4 triplet. There was a total of 10664 triplets (Monkey K: 3944, Monkey P=6720). The median split analysis is presented in three forms: 1) for an example triplet, 2) for all triplets, followed by averaging of V1-to-V4 GC, 3) for median splits based on the averaged 7A-to-V1 GC, followed by averaging of V1-to-V4 GC. For approaches 2) and 3), individual peak frequencies were aligned across monkeys. The median split analysis was repeated for the inverse direction, splitting epochs based on V1-to-V4 gamma-band GC jackknife replications (average of the peak ± 5 Hz), followed by averaging of 7A-to-V1 GC.

### Binning of sorted trials

To determine correlation values after sorting and binning the data, JR values of 7A-to-V1 beta-band GC were sorted and binned into either 5, 10, 50, or 100 bins of equal size. Per bin, JR values of 7A-to-V1 beta-band GC and of V1-to-V4 gamma GC were averaged. The Spearman rank correlation was determined across bins. When data were combined across monkeys, correlation coefficients were averaged across triplets for each monkey, and then averaged across monkeys. Statistical testing was based on permutations of bin order. Per permutation, bin order was randomized, and correlation coefficients were computed and averaged as for the observed data. This was applied 10000 times, resulting in minimal p-values of 0.0001.

### Jackknife Correlation

We aimed at quantifying the correlation between epoch-by-epoch fluctuations in two GC influences. This is normally precluded by the fact that GC influences are not defined per single data epoch (without substantially sacrificing spectral resolution and/or signal-to-noise ratio). Therefore, we used the Jackknife Correlation (JC), which quantifies the correlation by first calculating GC influences for all leave-one-out subsamples (i.e. the jackknife replications of all epochs) and then correlating these values (Richter et al., 2015). For each leave-one-out subsample, the GC or any other smooth function F of the data can be defined as follows:

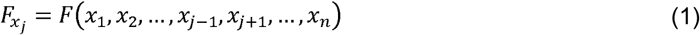

, where x specifies the pair of recording sites and j specifies the index of the left-out observation, here the epoch. Attend contra and attend ipsi conditions were combined for the JC analysis. The JC is defined using the following formula:

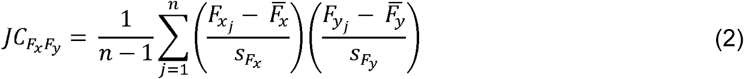

where *n* is defined as the number of jackknife replications and is equal to the total number of epochs, 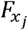 and 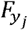 are the jackknife replications, 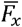 and 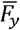 are the means of the jackknife replications, 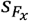 and 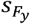 are the standard deviations of the jackknife replications. To use the JC with the Spearman correlation metric, we applied the above formula on the ranks of 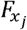 and 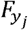.

For the statistical analysis of the observed JC values, we created a distribution of 1000 JC values under a realization of the null hypothesis of independence between the 7A-to-V1 and V1-to-V4 GC components of each triplet. This was realized by calculating JC between randomly permuted orderings of jackknife replications of 7A-to-V1 and V1-to-V4 GC influences, which is equivalent to calculating the JC between GC influences after leaving out a random epoch for the 7A-to-V1 GC and a random epoch for the V1-to-V4 GC influence without replacement. To control for multiple comparisons across the frequency-frequency combinations, the max-based approach (see above) was again employed. For each permutation, the maximum absolute Spearman’s rho value was selected, giving rise to an omnibus distribution of surrogate correlation coefficients. For the example triplet, the observed values were compared to the distribution of maximum absolute surrogate correlation values. For the average across triplets, the average was first calculated over triplets, then over monkeys, after aligning to their respective peak frequencies. For statistical testing, the same randomization was applied to all triplets of a given monkey, subsequent averaging was performed as for the observed data, and the maximum-based multiple comparison correction was applied as for the example triplet. An additional JC analysis first averaged all 7A-to-V1 site pairs and all V1-to-V4 site pairs to form one triplet per monkey. JC values were then averaged over monkeys. For statistical testing, the randomization was applied to this triplet per monkey, followed by averaging over monkeys, and the max-based multiple comparison correction.

For testing spatial specificity, we analyzed recording site triplets, which did not share the same V1 site (see Results for additional details): 7A-to-V1_a_V1_b_-to-V4. Since a vast number of such triplets exist, yet we wished to select a number equal to the original number of triplets to control potential statistical bias, we randomly selected a number of 7A-to-V1_a_V1_b_-to-V4 triplets that matched the original number of 7A-to-V1-to-V4 triplets evaluated for each monkey. We repeated this procedure 100 times and averaged the outcomes. Results were plotted against the average distance between the two V1 sites, V1_a_ and V1_b_, obtained for each distance interval.

### Weighted Jackknife Correlation

JCs might be particularly strong for triplets with strong 7A-to-V1 beta-band GC and/or strong V1-to-V4 gamma-band GC. To test for this, we weighted JC by multiplying it with the product of the 7A-to-V1 beta and the V1-to-V4 gamma GC magnitudes. Weights were normalized such that the average weight across triplets was one. The weighted and the unweighted JC values were separately averaged per monkey and then across monkeys. Statistics were based on random exchanges between weighted and unweighted values, within each monkey, and subsequent averaging as for the observed data. Randomization was applied 1000 times.

### Lagged Jackknife Correlation (LJC)

We were interested in whether the correlation between top-down beta GC and bottom-up gamma GC depended on their time lag. We started out by using the JC as described above. To smooth the results against shifts in peak frequencies over time, we used the average of the range of the top-down beta GC peak ±5 Hz, and the bottom-up gamma GC peak ±10 Hz. Since a given jackknife replication eliminated the same epoch for the calculation of both GC influences, this established the correlation at zero time-lag. To investigate the dependence of JC on time lag between GCs, we computed the JC between GC influences calculated from epochs that were offset by a variable lag. The epochs were stepped at intervals of t=0.005 s. The offsets were stepped at τ =0.005 s. Note that stepping of intervals and offsets was in principle independent and could have been different, but it was chosen to be identical to speed up computation. We refer to this as lagged JC (LJC):

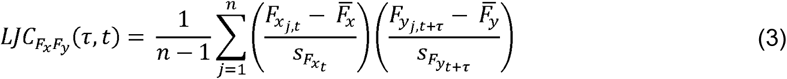

τ was chosen to cover a range of lags from -0.5 s to 0.5 s. The GC calculation itself was as in the previous zero-lag JC, using 0.5 s and the tapering specified above. LJC was calculated across trials, i.e. leaving out an entire trial at a time (this is different from the previous zero-lag JC, which used multiple non-overlapping epochs per trial if available). The data that was available per trial allowed for multiple realizations of the two epochs with a particular lag. For each lag, LJC was calculated separately for all possible realizations and averaged. The number of possible realizations decreases as the lag between top-down beta GC and bottom-up gamma GC increases, resulting in fewer LJC computations that are averaged. This results in a noisier estimate at larger lags, but no systematic bias in the average JC value. The number of epochs that each LJC is computed upon always equals the number of trials. Formally, this implementation of the LJC is defined as:

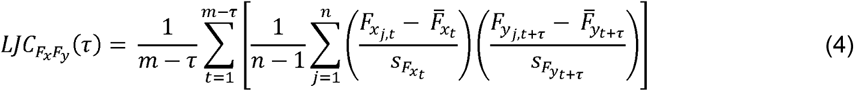

where m is the number of 0.5 s windows, stepped at 0.005 s, that fit into the trial length of 1.7 s.

Statistical significance was assessed using the same logic as used for the JC, where the epoch order of one member of the JC was permuted with respect to the other. For the LJC, the permutation was identical for each time step and lag, to be conservative. Multiple comparison correction must take place over the multiple lags, which was achieved by taking the maximum absolute Spearman’s rho value across lags for each permutation. The resulting distribution was used to assess the probability that the observed result at each lag occurred by chance. The observed and the permutation metrics were first averaged over all triplets per monkey and then averaged over the two monkeys, to give equal weight to both subjects.

We wished to assess whether the LJC peak lag of -0.105 s was significantly different from a lag of zero. We did so using a jackknife method to determine the standard error of the peak position in milliseconds (Efron, 1981). In this case, we leave out a specific triplet to assess the variability of the peak. The jackknife procedure causes a compression of the variance (Richter et al., 2015), thus the 0.005 s sampling grid would not be sufficient to represent the peak positions of the jackknife replications. To account for this, we cubic-spline interpolated each replication to a resolution of 1e-6 s, which proved adequate to represent the variance of the peak. The peak of each jackknife replication was found using a Gaussian fit of the smoothed correlation as a function of lag (findpeaksG.m by T.C O’Haver). We then derived the standard error of the estimator, and converted this to a t-score by dividing the mean peak lag value of the jackknife replications by the estimated standard error. The significance of this t-value was then assessed against Student’s t-distribution. At the group level, this procedure entails concatenating the data from both monkeys, and leaving out each triplet once. Based on this group estimate of the standard error, a t-value is derived, as above, and assessed for statistical significance.

## Results

### Top-Down Versus Bottom-Up Spectral Asymmetries and their Stimulus and Task Dependence

We performed electrocorticographic (ECoG) recordings from large parts of the left hemisphere in two macaque monkeys performing a selective attention task (Fig. 1). From the ECoG recording sites, we selected three ROIs (Fig. 1B), according to the following criteria: 1) One ROI should be a high-level control area, one ROI a low-level visual area, and one ROI a higher-level visual area; 2) ECoG coverage should be as good as possible; 3) ROIs should not be directly abutting, to avoid ambiguity in boundary definition and to minimize volume conduction. These criteria led to the selection of areas 7A, V1 and V4. The ROI pair 7A-V1 constitutes a clear top-down pathway with documented projections from a very high-level control area to primary visual cortex (Markov et al., 2014; Bastos et al., 2015c; Michalareas et al., 2016). The ROI pair V1-V4 constitutes a clear bottom-up pathway emerging from V1, i.e. the area targeted by the top-down 7A-to-V1 influence. For both ROI pairs, the ECoG provided good coverage. The central aim was to determine whether the top-down influence was modulating the bottom-up influence.

To assess the individual frequency bands for each monkey (monkey K and monkey P), we first computed the power spectra during visual stimulation. Area 7A shows strong beta-band peaks in both monkeys (Fig. 2E, F, monkey K: ≈17 Hz; monkey P: ≈13 Hz). Areas V1 and V4 show gamma frequency peaks (Fig. 2A-D, monkey K: ≈76 Hz; monkey P: ≈60 Hz). Beta activity is visible in V4 and V1 of both monkeys at their matching peak frequencies found in area 7A. In area V4 of both monkeys, there are distinct beta peaks. In area V1, monkey K shows a distinct beta peak, and monkey P shows a shoulder in the power spectrum, at the respective beta frequency.

We determined the dominant interareal communication frequencies for each monkey by calculating the pair-wise phase consistency (PPC), a frequency-resolved measure of synchronization (Vinck et al., 2010), between the V1-V4 and V1-7A ROI pairs (Fig. 2G-J). Gamma band synchronization was present for both ROI pairs in both monkeys with peaks at ≈76 Hz in monkey K and in a range of 58–65 Hz in monkey P. Beta peaks were present between both ROI pairs: at ≈17 Hz in monkey K and at ≈12 Hz in monkey P. For the further analyses, data from both monkeys were combined, by aligning their individual beta and gamma peaks ±10 Hz and averaging across monkeys.

When determining individual beta and gamma frequencies, we selected the dominant peaks in the respective frequency ranges. It has been shown that both beta and gamma frequencies show substantial inter-individual variability, which can largely be explained by genetic factors (Vogel, 1970; van Pelt et al., 2012). Note that while the beta peak in monkey P has its maximum at 12-13 Hz, at the border between the alpha and beta band, the peak is strongly asymmetric with a sharp rise to the maximal value and a slower falloff, such that most of the peak falls into the classical beta-frequency band. This observation, together with the fact that it corresponded phenomenologically to the clear beta peak in monkey K, led us to refer to it as beta, rather than alpha. The further analyses confirmed that the beta rhythms in both monkeys exerted qualitatively the same effects. Yet whether they reflect the same underlying physiological process cannot be determined on the basis of the ECoG recordings alone. Some of the power and PPC spectra showed also a theta-band peak, which is not further investigated, because the focus of this study is on the interaction between beta and gamma rhythms.

To demonstrate that interareal gamma-band synchronization is stimulus driven (Bosman et al., 2012; Grothe et al., 2012), we contrasted PPC between the fixation and stimulation conditions. Figure 3A, B shows significantly enhanced gamma-band synchronization between ROI pairs V1-V4 and 7A-V1 once the stimulus has appeared, in contrast to an almost flat spectrum when no stimulus is present. This finding is consistent with gamma-band oscillations occurring as a result of stimulus drive. In contrast, beta-band synchronization for both ROI pairs is present already during the pre-stimulus fixation period, suggesting an endogenous origin (Fig. 3A, B). Beta synchronization is maintained during the stimulation period, consistent with an ongoing top-down influence.

We next assessed the dominant directionality of interareal synchronization and its attentional modulation. We quantified directionality of synchronization by means of Granger causality (GC) (Granger, 1969; Ding et al., 2006; Bressler and Seth, 2011). As shown by Bastos et al. (2015c), and extended to humans by Michalareas et al. (2016), the top-down beta-band influence of area 7A to V1 is significantly greater than the bottom-up beta-band influence of V1 to 7A (Fig. 3E). This top-down beta-band influence is significantly increased when attention is directed to the visual hemifield contralateral to the recording grid (Fig. 3C), consistent with an earlier report (Bastos et al., 2015c). Between V1 and V4, the gamma-band influence is stronger in the bottom-up direction from area V1 to V4 (Fig. 3F). The bottom-up gamma-band influence of V1 to V4 was significantly increased with attention (Fig. 3D).

Note that the beta-band GC influence between V1 and V4 was stronger in the bottom-up than top-down direction (Fig. 3F), in contrast to what has been reported for the same dataset (Bastos et al., 2015c). This is due to differences in preprocessing between the present and the previous study. It shows that for this particular pair of areas, the beta-band directionality is not robustly determined by their hierarchical relationship. However, the difference in preprocessing did not affect other area pairs, leaving the overall pattern intact, in which gamma-band influences are stronger in the bottom-up direction and beta-band influences are stronger in the top-down direction. This overall pattern has also been confirmed in an independent dataset from 43 human subjects (Michalareas et al., 2016), and in this large human dataset, the beta-band influence between V1 and V4 also does not show an unequivocal directionality, despite the overall pattern across all area pairs. Note that both in the human MEG data, and in the macaque ECoG data presented here, V1-to-V4 GC did not significantly exceed V4-to-V1 GC for all frequencies, that is, there was not a general broadband shift.

## Top-Down Beta-Band Influences Correlate with Bottom-Up Gamma-Band Influences

Spontaneous endogenous increases in 7A-to-V1 beta GC may lead to increases in stimulus-driven V1-to-V4 gamma GC. A resulting correlation between epoch-by-epoch fluctuations in the two GC influences is difficult to quantify, because GC influences are not defined per single data epoch (without substantially sacrificing spectral resolution and/or signal-to-noise ratio). To surmount this problem, we used a jackknife approach. A jackknife replication (JR) of all epochs consists of all epochs except one, that is, an all-but-one subsample of the epochs. There are as many JRs as there are epochs, because each epoch can be left out once. If the GC in a given JR is high, this reflects a low GC in the left-out epoch, and vice versa. This approach allows to median split epochs according to GC, and to sort and bin epochs according to GC. Ultimately, it also allows to calculate the correlation between epoch-wise top-down and bottom-up GC, a technique that we recently described and named Jackknife Correlation (JC) (Richter et al., 2015).

We first investigate this for one example triplet of recording sites from monkey K. Data epochs were median split based on the GC JRs. When we select the epochs in which V1-to-V4 gamma GC is weak, 7A-to-V1 beta GC is almost absent; by contrast, when we select the epochs in which V1-to-V4 gamma GC is strong, 7A-to-V1 beta GC shows a pronounced peak (Fig.4A, left panel). A similar dependency exists in the other direction. When we select epochs with weak 7A-to-V1 beta GC, V1-to-V4 gamma GC is small; when we select epochs with strong 7A-to-V1 beta GC, V1-to-V4 gamma GC is much larger (Fig. 4A, right panel). These results indicate a relationship between 7A-to-V1 beta GC and V1-to-V4 gamma GC.

**Figure 4.**
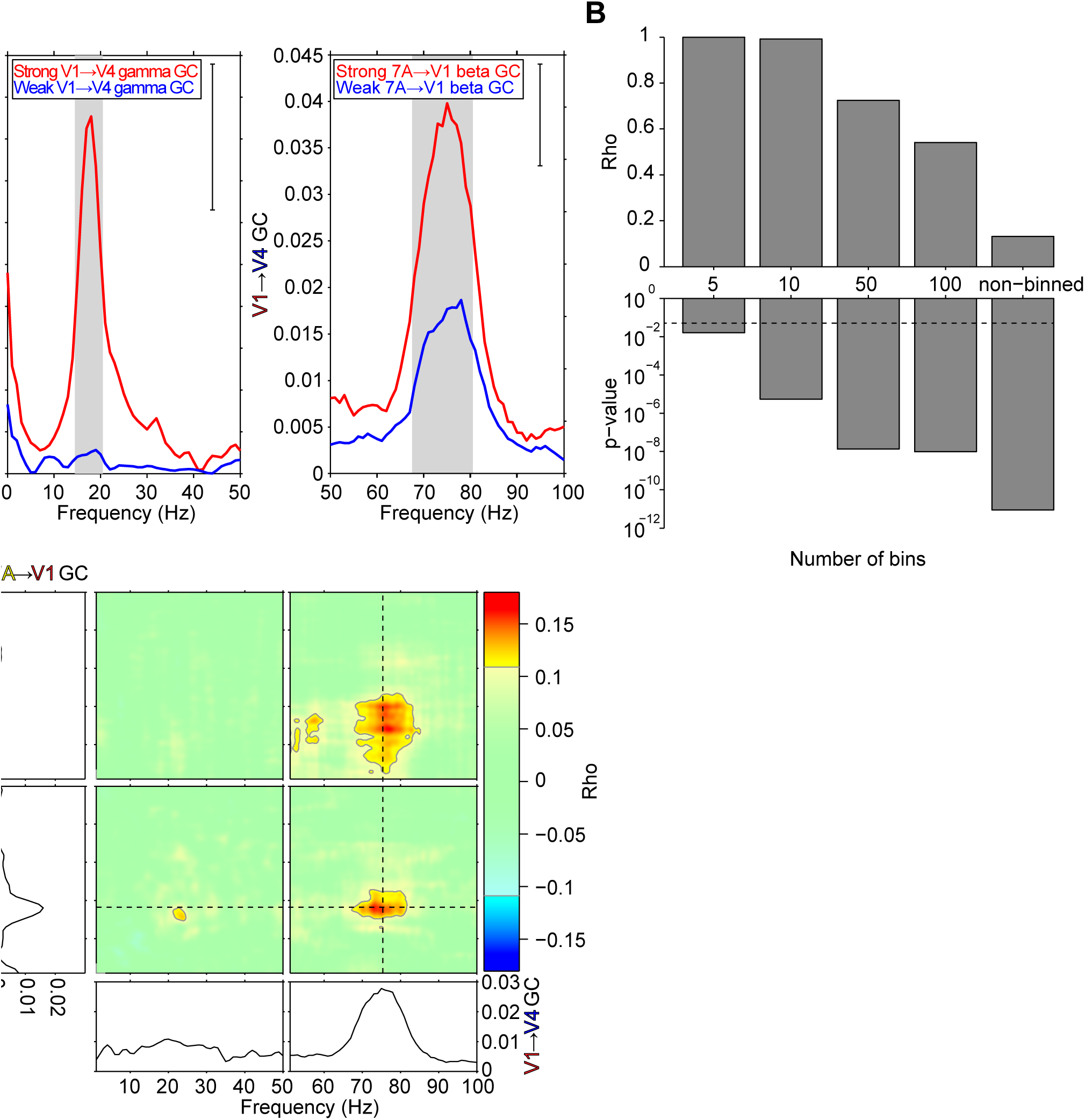
Example triplet: Median split, correlation across binned epochs, and JC. ***A***, left panel: 7A-to-V1 GC for epochs median split by the V1-to-V4 gamma GC jackknife replications; right panel: V1-to-V4 GC for epochs median split by the 7A-to-V1 beta GC jackknife replications. Gray background shading indicates significant differences (p<0.05, two-tailed non-parametric randomization test, corrected for multiple comparisons across frequencies). Inset brackets denote the minimum separation required for significance. ***B***, The correlation between 7A-to-V1 beta GC and V1-to-V4 gamma GC for the sorted data divided into 5, 10, 50, or 100 bins, and without binning. Corresponding p-values for each correlation coefficient are shown below with a dashed line marking p=0.05. ***C***, The 4 colored panels show JC for the selected 7A-to-V1-to-V4 triplet. The frequencies of 7A-to-V1 GC are shown on the vertical axis, the frequencies of V1-to-V4 GC are shown on the horizontal axis. The frequency ranges 1-50 Hz and 51-100 Hz are shown separately, because they required slightly different spectral analyses (see Materials and Methods). Non-significant regions are partially masked by white (p<0.001, two-tailed non-parametric randomization test, corrected for multiple comparisons across both frequency axes). The line plots at the bottom show the GC spectrum for the corresponding V1-to-V4 site pair. The line plots to the left show the GC spectrum for the corresponding 7A-to-V1 site pair. Dashed lines mark the top-down beta GC spectral peak and the bottom-up gamma GC spectral peak.

We next asked whether this relationship holds when we move from the coarse median split to finer and finer bins, and calculate the Spearman rank correlation across bins. We sorted epochs according to 7A-to-V1 beta GC JRs and binned them into 5, 10, 50 or 100 bins. Per bin, the 7A-to-V1 beta GC JRs and the V1-to-V4 gamma GC JRs were averaged, and the correlation between them was calculated across bins. This analysis revealed that the relationship found for the median split indeed held for finer bins (Fig. 4B). Finally, we used JC to base the correlation analysis on single epochs, which confirmed the relationship even at this most fine-grained level (Fig. 4B, rightmost bars). The finer the binning, the lower the correlation value, and the lower the p-value, that is, the more significant the correlation. This dependence of correlation and p-value on bin size has been previously described as a general consequence of binning (Richter et al., 2015). Essentially, the widely-used sorting-and-binning approach, through averaging observations per bin and thereby removing noise, leads to dramatic increases in correlation values; yet this comes at the expense of statistical power and it precludes an inference on the epoch-by-epoch correlation, which requires the JC approach.

To investigate the frequency-specificity of this correlation, JC was calculated between all possible combinations of top-down and bottom-up frequencies, both ranging from 1-100 Hz. This analysis revealed a correlation between 7A-to-V1 GC and V1-to-V4 GC in the beta band and in the gamma band (Fig. 4C, lower left and upper right quadrants). Critically, and further confirming the median-split and the sorting-and-binning analysis, 7A-to-V1 beta GC shows a significant positive correlation with V1-to-V4 gamma GC (Fig. 4C, lower right quadrant). The peak of this cross-frequency interaction is well aligned with the 7A-to-V1 beta and V1-to-V4 gamma GC peak frequencies (Figure 4C, dashed lines). There is no significant JC between 7A-to-V1 gamma and V1-to-V4 beta GC, even though 7A-to-V1 gamma GC is significantly correlated to V1-to-V4 gamma GC, and 7A-to-V1 beta GC is significantly correlated to V1-to-V4 beta GC.

We then repeated these analyses for all possible triplets (Fig. 5). Figure 5A, B, C uses the same format as Figure 4A, B, C, but shows the grand average over all triplets and the two monkeys after alignment by their respective top-down beta, and bottom-up gamma peak frequencies. The pattern of results found for the example triplet held in the grand average, even though average effect size was smaller. Figure 5D shows the probability distribution across triplets of JC values between 7A-to-V1 beta GC and V1-to-V4 gamma GC, averaged over monkeys. The distribution shows a positive skew and greater mass above zero, indicating a greater number and magnitude of positive correlations, though with a substantial number of negative correlations, which partially accounts for the low average JC value.

**Figure 5.**
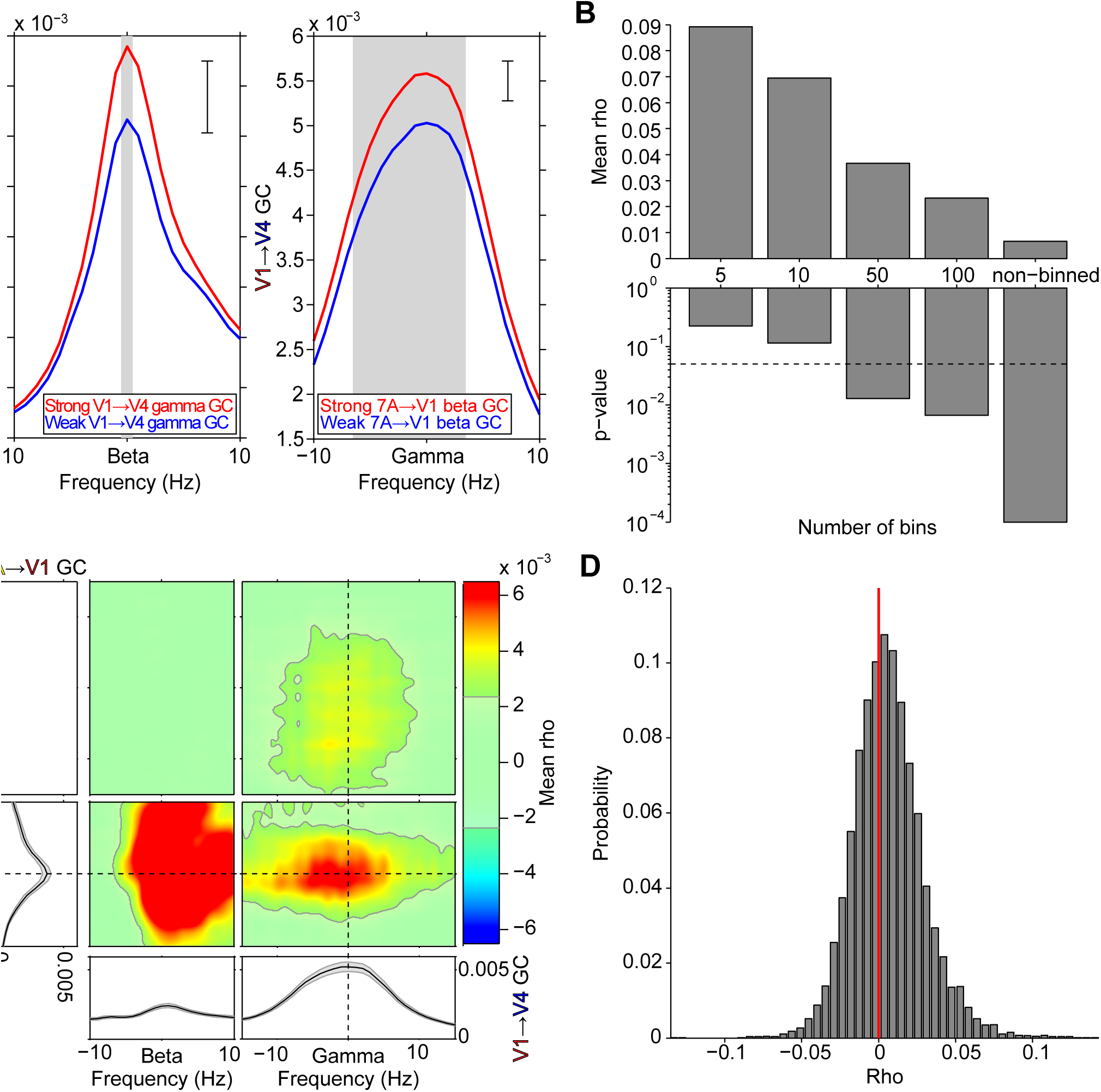
Average over all triplets and both monkeys: Median split, correlation across binned epochs, and JC. (***A***, ***B***, ***C***) Same format as Figure 4A, B, C, but averaging over all triplets and both monkeys, after aligning to their individual beta and gamma peak frequencies. ***D***, Probability distribution across triplets of JC values between 7A-to-V1 beta GC and V1-to-V4 gamma GC, averaged over monkeys.

To determine whether greater GC values resulted in greater JC values, we weighted the JC value of each triplet by the product of the respective triplet’s top-down beta and bottom-up gamma GC values, followed by averaging over triplets. This resulted in an increase of the mean JC value from 0.006 to 0.014, that is, a ≈230% increase (p<0.001, two-tailed non-parametric randomization test). This indicates that site pairs with larger GC magnitudes, which are less susceptible to the influence of noise, give rise to higher correlation coefficients.

Next, we tested whether the correlation was affected by attentional state. We calculated the JC between 7A-to-V1 beta GC and V1-to-V4 gamma GC, separately for the attend-contra and attend-ipsi conditions. The JC was significant for each attention condition separately (p<0.001). Selective attention to the right hemifield stimulus, activating the recorded left hemisphere, enhanced JC values by 17% (attend-contra: 0.007; attend-ipsi: 0.006, p=0.006, two-tailed paired non-parametric randomization test). This effect is likely related to the observed increase in JC, when triplets are weighted by GC values, because attention increased both 7A-to-V1 beta GC and V1-to-V4 gamma GC (Fig. 3 C, D).

The sorting-and-binning analysis showed that the JC increases substantially when 7A-to-V1 GC and V1-to-V4 GC are averaged per bin, thereby removing noise across individual triplets within bins. The bin-wise averaging precludes an inference on the epoch-by-epoch correlation. Therefore, we explored an alternative approach to reduce noise, while retaining an epoch-by-epoch correlation. Per epoch, we averaged GC JRs over all 7A-to-V1 site pairs, and we averaged GC JRs over all V1-to-V4 site pairs. Thus, per monkey, we constructed one triplet comprising the average 7A-to-V1 GC and the average V1-to-V4 GC. Figure 6A, B, C shows the resulting analyses in the same format as the respective panels in Figures 4 and 5. After reduction of noise across site pairs, the pattern of results was confirmed, yet with substantially increased correlation values. Importantly, this also holds for the epoch-wise JC values shown in the rightmost bars of Figure 6B and in Figure 6C. The JC value between 7A-to-V1 beta GC and V1-to-V4 gamma GC increased from 0.006 in the average across individual triplets to 0.12, that is, it showed a 20-fold increase. This is a striking demonstration of the utility of spatial averaging for exposing the epoch-wise correlation. When we additionally give up the epoch-wise calculation and smooth the GC values further through binning of epochs, correlations increase even more and exceed values of 0.6 for a typical 10-bin scheme (Fig. 6 B).

**Figure 6.**
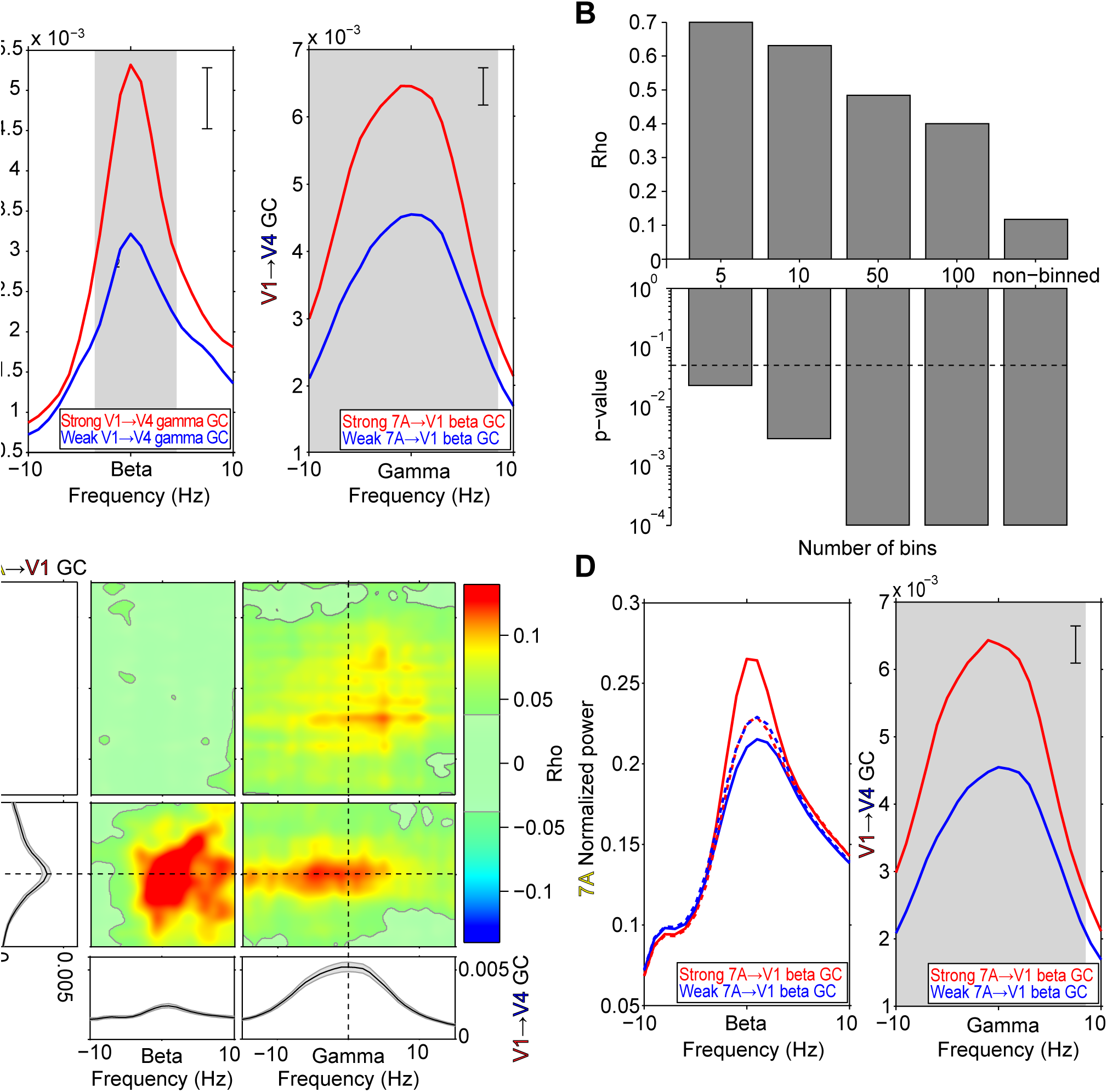
Triplet based on average 7A-to-V1 GC and average V1-to-V4 GC: Median split, correlation across binned epochs, and JC. (***A***, ***B***, ***C***) Same format as Figure 5A, B, C, but for a triplet formed by the average 7A-to-V1 GC jackknife replications and the average V1-to-V4 GC jackknife replications, per monkey, and then averaged over monkeys. ***D***, Left panel: 7A beta-band power spectra after median split by 7A-to-V1 beta GC jackknife replications. Solid lines: Before stratification; Dashed lines: After stratification. Right panel: Same as right panel of ***A***, but after stratification for 7A beta power.

To investigate whether V1-to-V4 gamma GC is truly dependent on 7A-to-V1 beta GC influences, rather than merely on 7A beta power, we stratified for beta power. We applied this to the median split analysis, because it lends itself best to stratification. As before, we median split epochs based on 7Ato-V1 beta GC JRs. The 7A power spectra aligned to the monkeys’ individual beta peak frequencies showed that stronger 7A-to-V1 beta GC was in fact related to higher 7A beta power (Fig. 6D, left panel, solid lines). This power difference is fully in line with the hypothesis that the top-down beta-band influence is generated in 7A. Power differences were sufficiently small, such that stratification rendered the power spectra almost identical (Fig. 6D, left panel, dashed lines). After stratification, V1-to-V4 gamma-band GC values remained almost unchanged compared to the values before stratification (compare right panels of Fig. 6A and D). Thus, V1-to-V4 gamma GC depended not on 7A power differences, but rather on actual 7A-to-V1 GC influences through synchronization.

### Spatial Resolution of the Correlation between Top-Down and Bottom-Up Influences

We next investigated whether the JC between 7A-to-V1 beta GC and V1-to-V4 gamma GC depended on involving the same V1 site, which would demonstrate spatial specificity at the level of recording sites. We tested this spatial specificity by pairing 7A-to-V1 beta GC to a specific V1 site, with V1-to-V4 gamma GC from a different V1 site. The distance that separated the two V1 sites was parametrically varied. For each V1 site, 5 sets of other V1 sites were defined that fell into pre-specified distance intervals (1 cm per interval, stepped by 2.5 mm, between 0 and 2 cm). Figure 7A shows one example V1 site (arrow) and illustrates with 5 colored lines the five distance intervals (colored lines were slightly displaced for illustration purposes). The average distance for each distance interval is marked with a filled circle. Figure 7B shows the resulting JCs, averaged over triplets and monkeys, as a function of distance. It can be seen that as the distance between the two V1 sites increases, there is a monotonic falloff of the correlation coefficient between 7A-to-V1 beta and V1-to-V4 gamma GC. This indicates that the physiological process linking 7A-to-V1 beta GC to V1-to-V4 gamma GC is not global, but rather spatially specific. As a consequence, any spatially non-specific fluctuations, for example of neuromodulators reflecting arousal fluctuations, are unlikely to cause the observed correlation.

**Figure 7.**
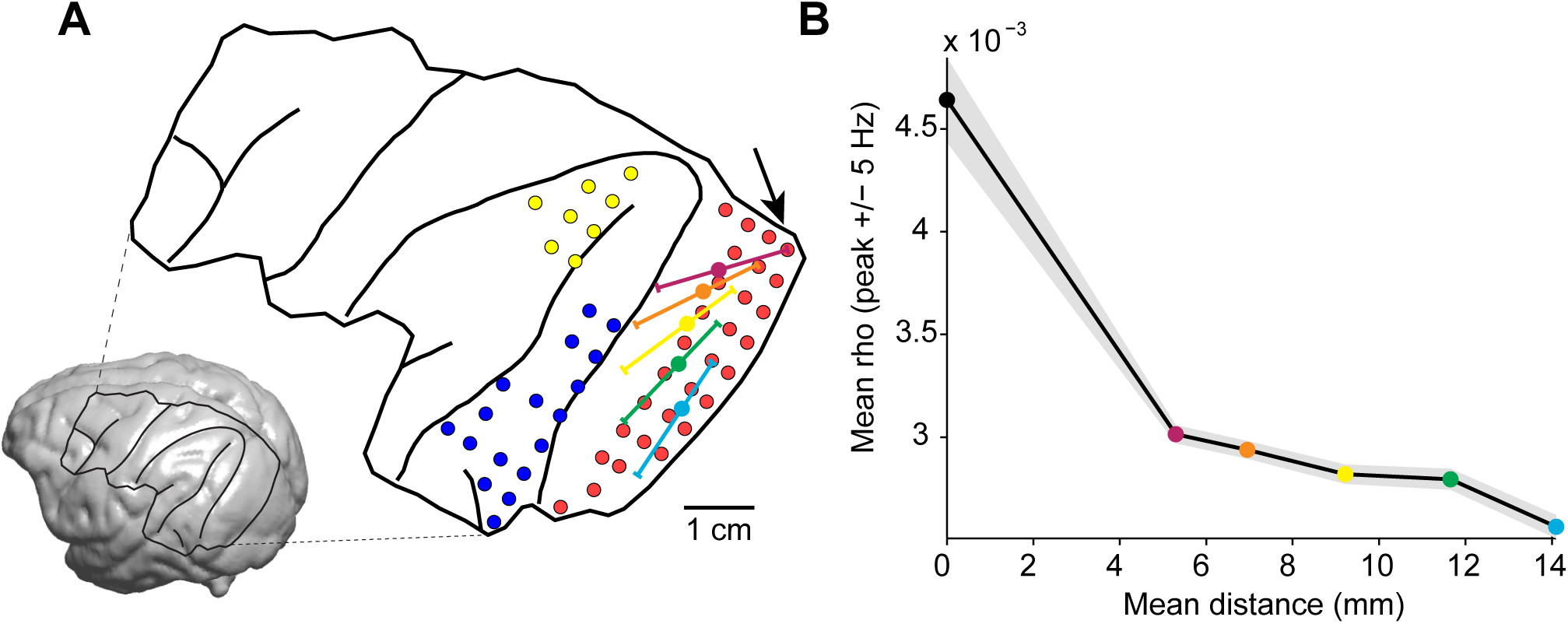
Spatial specificity of correlation between top-down beta and bottom-up gamma GC. ***A***, Inset: MRI of monkey K with the surgical trepanation and prominent sulci overlaid. The blow-up shows recording sites in 7A (yellow), V1 (red), and V4 (blue) that were analyzed. The V1 site marked by the arrow serves as an example to demonstrate the distance intervals from which another V1 site may have been chosen, which were 0 to 1 cm (magenta), 0.25 to 1.25 cm (orange), 0.5 to 1.5 cm (yellow), 0.75 to 1.75 cm (green), and 1 to 2 cm (cyan). Solid circles mark the average distance between V1 sites that were chosen for each distance interval. The particular arrangement of the distance intervals was chosen solely for the purpose of illustration. ***B***, Average jackknife correlation between 7A-to-V1 beta GC and V1-to-V4 gamma GC, averaged over triplets, then over monkeys. The solid black circle shows the result for standard triplets, that is, triplets where the 7A-to-V1 beta GC and the V1-to-V4 gamma GC are connected through the same V1 site. As the distance between V1 sites increases, there is a monotonic falloff in the correlation coefficient. The mean distance between V1 sites is color-coded as solid circles matching the mean distances of the five distance intervals shown in ***A***. The error region shows ±1 SEM across triplets per monkey, then averaged over monkeys.

### Top-down Beta Leads Bottom-up Gamma in Time

We have established that spontaneous fluctuations in endogenous top-down beta GC are correlated with fluctuations in stimulus-driven bottom-up gamma GC. To investigate whether the data contain evidence in support of a causal relation, we assessed whether top-down beta GC is predictive of subsequent bottom-up gamma GC. To accomplish this, we extended the JC by adding a temporal dimension, similar to time-lagged cross-correlation. We compute the JC on time-frequency data, where we systematically offset 7A-to-V1 beta GC JRs from V1-to-V4 gamma GC JRs by positive or negative lags. We call this procedure lagged jackknife correlation (LJC). This quantifies at what time delay between the top-down beta-band influence and the bottom-up gamma-band influence the JC between them is largest. We computed LJC for each triplet. The red curve in Figure 8 shows the LJC averaged over triplets and monkeys and exhibits a peak at -0.105 s, indicating that 7A-to-V1 beta GC led V1-to-V4 gamma GC by 0.105 s (t(10663) = -7.576, p<<0.001, two-tailed jackknife-based t-test). The LJC dropped off slowly, which likely reflects slow dynamics of the underlying GC influences together with the fact that the windows used for GC influence estimation result in low-pass filtering of GC influence time courses. For comparison, we performed the same LJC analysis between 7A-to-V1 gamma GC and V1-to-V4 beta GC. This combination had not shown a significant correlation in the non-lagged case (Fig. 5C, upper left quadrant), which was confirmed for the lagged case (Fig. 8, gray line).

**Figure 8.**
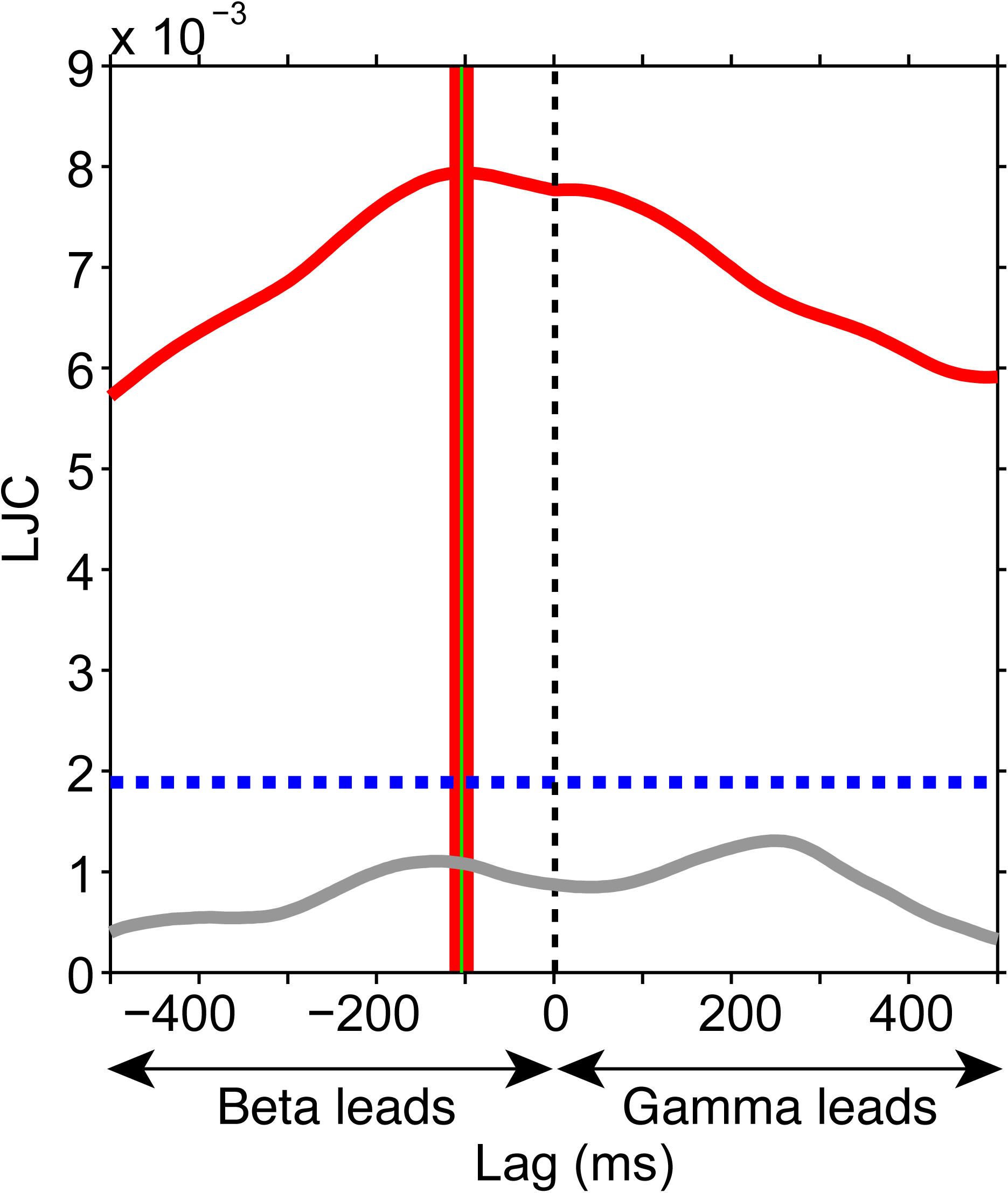
Lagged jackknife correlation (LJC) analysis. ***A***, The solid red curve shows the LJC between 7A-to-V1 beta GC and V1-to-V4 gamma GC, averaged over triplets, then over monkeys. The blue dashed line denotes the significance threshold (p=0.05, two-tailed non-parametric randomization test, corrected for multiple comparisons across lags). The green vertical line indicates the lag of the maximum LJC value, with ±1 SEM indicated in red. The lag of the peak LJC value was significantly different from zero (t(10663) = -7.576, p<<0.001, two-tailed). Solid gray curve: LJC between 7A-to-V1 gamma GC and V1-to-V4 beta GC, averaged over triplets, then over monkeys, which showed no significant peaks.

## Discussion

We used LFP recordings from 252-channel ECoG arrays covering large parts of the left hemispheres of two macaques to analyze the interaction between top-down and bottom-up influences, both quantified by Granger-causality (GC). Top-down influences were assessed between area 7a at the top of the visual hierarchy and V1 at the bottom. Bottom-up influences were assessed between V1 and V4, a known feedforward pathway carrying stimulus driven input. 7A-to-V1 GC showed a beta-band peak, which did not require visual stimulation and thus was endogenously generated, which was significantly larger in the 7A-to-V1 than the V1-to-7A direction, and which increased with selective attention. V1-to-V4 GC showed a gamma-band peak, which was stimulus driven, which was significantly larger in the bottom-up than the top-down direction, and which also increased with selective attention. Jackknife Correlation between top-down beta-band influences and bottom-up gamma-band influences revealed a positive cross-frequency interaction. This interaction was spatially specific, as it was maximal between top-down and bottom-up interareal influences that shared the same V1 site. Finally, top-down beta-band influences best predicted bottom-up gamma-band influences ∼0.1 s later, suggesting that the cross-frequency interaction is causal. Therefore, we conclude that 7A-to-V1 beta-band influences enhance V1-to-V4 gamma-band influences.

Noise can affect GC estimates (Nalatore et al., 2007; Vinck et al., 2015). Fluctuating shared noise could in principle generate correlation between GC fluctuations. Yet, a predominant source of shared noise, volume conduction, is strongly attenuated in our signals due to bipolar derivation (Trongnetrpunya et al., 2015). Furthermore, influences of shared noise on the two GC metrics should typically occur nearly simultaneously and therefore, the lagged JC would peak at zero-lag, whereas we found a significant lag of ∼0.1 s. Also, local noise can affect GC estimates, yet these effects do not necessarily explain a positive correlation between GC influences. For example, enhanced noise in a recording site can lead to an overestimation of the GC influence onto that site, and at the same time to an underestimation of the GC influence of that site onto other sites (Bastos and Schoffelen, 2015a). If this applied to our V1 recording sites, fluctuations in their noise level would artefactually reduce the observed correlation and thereby lead to an underestimation of the true correlation value. Finally, the observed increase in average JC after weighting site triplets by the GC values of their constituent site pairs argues against a confounding role of noise. The effect of noise on GC is larger for weaker GC. Thus, if noise had generated the observed JC, more weight for strong-GC triplets would have reduced the average JC. By contrast, we observed an increased average JC, which is consistent with our interpretation of actual top-down influences affecting actual bottom-up influences.

Additional alternative scenarios concern physiological modulatory effects that act independently on both, 7A-to-V1 beta GC and V1-to-V4 gamma GC, without a direct causal link between the two GC influences. Those interareal GC influences can probably only be affected at the circuits containing the respective projection neurons and at the circuits containing their postsynaptic target neurons, that is, not at the corresponding interareal axonal projections. Thus, to explain our observations, such modulatory effects would need to modify 7A circuits to enhance their beta outflow, and 0.1 s thereafter modify V1 circuits to enhance their gamma outflow. Alternatively, they would need to modify V1 circuits to enhance their beta susceptibility, and 0.1 s thereafter modify V4 circuits to enhance their gamma susceptibility. Additional mechanisms would be necessary to explain the observed spatial specificity. Thus, these alternative interpretations require complex sets of assumptions. Certainly, widespread or even global neuromodulatory fluctuations, as potentially associated with arousal fluctuations, cannot explain the observed pattern of results.

The most parsimonious interpretation seems to be the following: The 7A-to-V1 beta-band influence onto a given V1 site enhances with 0.1 s delay that site’s V1-to-V4 gamma influence. We propose that this cross-frequency interaction constitutes a mechanism for spatially selective attention. If correct, this entails that the top-down control of selective attention corresponds fully or partly to top-down beta-band influences, and the ensuing preferential bottom-up routing of the attentionally selected stimulus corresponds fully or partly to bottom-up gamma-band influences. Thus, epoch-by-epoch fluctuations in spatially selective attention could well generate the observed correlations. In fact, we would like to identify the epoch-by-epoch GC fluctuations that are the basis for the observed correlations with epoch-by-epoch fluctuations of spatially selective attention. Several recent studies have revealed that attention samples stimuli rhyhthmically, with a predominant sampling rhythm in the theta and/or alpha range (7-13 Hz), which is multiplexed across the sampled stimuli (Landau and Fries, 2012; Fiebelkorn et al., 2013; Holcombe and Chen, 2013; Fries, 2015; Landau et al., 2015; VanRullen, 2016). Future studies will need to investigate whether the correlations described here are specifically driven by those attentional sampling rhythms.

Numerous studies in visual cortex have reported gamma-band synchronization within and between visual areas (Gray and Singer, 1989; Engel et al., 1991; Kreiter and Singer, 1996; Tallon-Baudry et al., 1996; Fries et al., 1997; Fries, 2001; Bichot et al., 2005; Taylor et al., 2005; Hoogenboom et al., 2006; Womelsdorf et al., 2006; Wyart and Tallon-Baudry, 2008), and numerous studies in parietal cortex have reported beta-band synchronization within parietal areas and between parietal and frontal areas (Buschman and Miller, 2007; Salazar et al., 2012; Dotson et al., 2014; Stetson and Andersen, 2014). Recent ECoG recordings covering both visual and parietal areas revealed that interareal beta-band influences predominate in the top-down and interareal gamma-band influences predominate in the bottom-up direction (Bastos et al., 2015c). These findings link parietal beta-band activity with visual gamma-band activity and suggest a concrete case of cross-frequency interaction (Bressler and Richter, 2015). In the present paper, we have tested some of the resulting predictions and found direct experimental support for such a cross-frequency interaction that allows top-down beta-band influences to enhance bottom-up gamma-band influences.

Cortical anatomy has revealed a distinct laminar pattern of top-down and bottom-up projections (Felleman and Van Essen, 1991; Markov et al., 2014). Bottom-up projections originate predominantly in superficial layers, and this predominance increases with the number of hierarchical levels bridged by the bottom-up projection. Note that an additional bottom-up pathway via the pulvinar originates in layer 5, and it might mediate the observed V1-to-V4 GC influences in the beta band (Shipp, 2003; Sherman, 2007; Schmid et al., 2012). Furthermore, bottom-up projections terminate predominantly in layer 4. Top-down projections originate predominantly in deep layers, and this predominance increases with the number of hierarchical levels bridged by the top-down projection. Furthermore, top-down projections terminate predominantly outside layer 4, primarily in layers 1 and 6. Determining how the respective top-down influences interact with local processing, and thereby ultimately with bottom-up influences, remains a central neuroscientific quest. One potential mechanism has been proposed in a model that entails details of both layer-specific anatomy and cellular biophysics (Lee et al., 2013), and that replicates the effect of top-down selective attention on bottom-up gamma-band coherence. The model implicates a subclass of inhibitory interneurons, the slow-inhibitory (SI) interneurons, as targets of top-down modulation. These cells may span multiple cortical laminae and thus are suitably situated for integration of neuronal activity across layers. A subpopulation of these cells, low-threshold spiking (LTS) cells, are found in deep layers of the cortex. In the model, LTS cells: 1) are hypothesized to receive top-down input, 2) are implicated in the generation of beta oscillations and in a resonant response to beta-rhythmic top-down input, 3) selectively modulate gamma-band activation in layer 2/3, leading to an enhanced gamma band output. Our present analysis confirms the central prediction of the Lee et al. (2013) paper, namely that specifically top-down beta-band influences enhance stimulus-driven gamma-band processes. Lee et al. show how this mechanism can support the implementation of attentional stimulus selection. The current results, which mechanistically link the previously reported attentional enhancements of top-down beta and bottom-up gamma influences, provide the hitherto missing experimental bridge. Together, experiments, modeling and model-testing data analysis have led to an intriguingly coherent understanding of the neuronal processes behind the implementation of attentional stimulus selection.

## Acknowledgements

This work was supported by DFG (SPP 1665, FOR 1847, FR2557/5-1-CORNET), EU (HEALTH-F2-2008-200728-BrainSynch, FP7-604102-HBP), a Europn Young Investigator Award, National Institutes of Health (1U54MH091657-WU-Minn-Consortium-HCP), the LOEWE program (NeFF) and the FLAG-ERA JTC 2015 project CANON (co-funded by NWO to CAB). We would like to thank Julien Vezoli for his assistance in ROI definition, co-registration and construction of anatomical renderings.

